# Brain stimulation boosts perceptual learning by altering sensory GABAergic plasticity and functional connectivity

**DOI:** 10.1101/2021.09.13.459793

**Authors:** Vasilis M Karlaftis, Polytimi Frangou, Cameron Higgins, Diego Vidaurre, Joseph J Ziminski, Charlotte J Stagg, Uzay E Emir, Zoe Kourtzi

## Abstract

Interpreting cluttered scenes —a key skill for successfully interacting with our environment— relies on our ability to select relevant sensory signals while filtering out noise. Training is known to improve our ability to make these perceptual judgements by altering local processing in sensory brain areas. Yet, the brain-wide network mechanisms that mediate our ability for perceptual learning remain largely unknown. Here, we combine transcranial direct current stimulation (tDCS) with multi-modal brain measures to modulate cortical excitability during training on a signal-in-noise task (i.e. detection of visual patterns in noise) and test directly the link between processing in visual cortex and its interactions with decision-related areas (i.e. posterior parietal cortex). We test whether brain stimulation alters inhibitory processing in visual cortex, as measured by magnetic resonance spectroscopy (MRS) of GABA and functional connectivity between visual and posterior parietal cortex, as measured by resting state functional magnetic resonance imaging (rs-fMRI). We show that anodal tDCS during training results in faster learning and decreased GABA+ during training, before these changes occur for training without stimulation (i.e. sham). Further, anodal tDCS decreases occipito-parietal interactions and time-varying connectivity across the visual cortex. Our findings demonstrate that tDCS boosts learning by accelerating visual GABAergic plasticity and altering interactions between visual and decision-related areas, suggesting that training optimises gain control mechanisms (i.e. GABAergic inhibition) and functional inter-areal interactions to support perceptual learning.

## Introduction

Interacting successfully in our environments entails discerning relevant information from clutter and identifying target objects in busy scenes. Training is shown to improve such perceptual judgements —a skill known as perceptual learning— by altering processing in sensory (i.e. visual) and decision-related (i.e. posterior parietal) areas. For example, perceptual learning (i.e. training on a visual discrimination task) has been shown to alter functional connectivity —as measured by rs-fMRI— between visual and posterior parietal cortex (Lewis et al., 2009). Further, we have previously shown that training in visual discrimination tasks alters GABAergic processing in visual cortex (Frangou et al., 2019, 2018) —as measured by MRS—, consistent with the role of GABAergic inhibition in brain plasticity (for a review see (Ip and Bridge, 2021)). Yet, the interactions between GABAergic plasticity in sensory areas and learning-dependent functional connectivity between sensory and decision-related areas for perceptual learning remain largely unknown.

Previous work has proposed that GABAergic inhibition shapes network connectivity (Kapogiannis et al., 2013; Mann and Paulsen, 2007; Shmuel and Leopold, 2008; Stagg et al., 2014). Here, we employ tDCS to modulate cortical excitability and test directly the link between local inhibitory processing in visual cortex —as measured by MRS GABA— and interactions between visual and decision-related areas (i.e. posterior parietal cortex) —as measured by static and time-varying (using Hidden Markov Models (HMM; (Vidaurre et al., 2018, 2017)) rs-fMRI connectivity. Anodal tDCS has been shown to be excitatory (Antal et al., 2004a; Nitsche and Paulus, 2000), result in decreased GABA levels in visual (Barron et al., 2016), frontal (Harris et al., 2019) and motor areas (Stagg et al., 2009), and facilitate visual (Frangou et al., 2018; Sczesny-Kaiser et al., 2016; Van Meel et al., 2016) and motor learning (O’Shea et al., 2017; Stagg et al., 2011). Further, anodal tDCS in the motor cortex has been shown to facilitate learning by decreasing local GABA levels and increasing functional connectivity within the motor network at rest (Bachtiar et al., 2015; Stagg et al., 2014).

We ask whether anodal tDCS in occipito-temporal cortex (OCT) facilitates learning and alters GABAergic processing and brain network interactions. We trained participants in a signal-in-noise discrimination task (i.e. participants were asked to detect radial vs. concentric patterns embedded in noise) that has been shown to involve occipito-temporal and posterior parietal cortex (Chang et al., 2014; Frangou et al., 2019, 2018; Mayhew et al., 2012). We tested for changes in task performance, MRS GABA+ and rs-fMRI connectivity in three groups of participants: two intervention groups who received anodal vs. sham tDCS during training on the task, a no-intervention group who received neither stimulation nor training in the task. Our results show that anodal OCT stimulation results in faster learning and decreased GABA+ during training, before these changes occur in the sham stimulation group. Further, anodal tDCS induces changes in occipito-parietal interactions and time-varying connectivity across the visual cortex. Finally, enhanced local temporal coherence in the visual cortex and decreased occipito-parietal connectivity relate to decreased OCT GABA+. Our findings suggest that tDCS boosts learning by altering visual GABAergic processing and network interactions between visual and decision-related areas.

## Results

### Anodal tDCS improves performance in signal-in-noise discrimination

We trained two intervention groups (anodal vs. sham tDCS on OCT) on a signal-in-noise (SN) task that involves participants discriminating shapes (radial vs. concentric Glass patterns) embedded in noise (Figure 1). Participants were asked to judge whether each stimulus presented per trial was radial vs. concentric.

**Figure 1.**
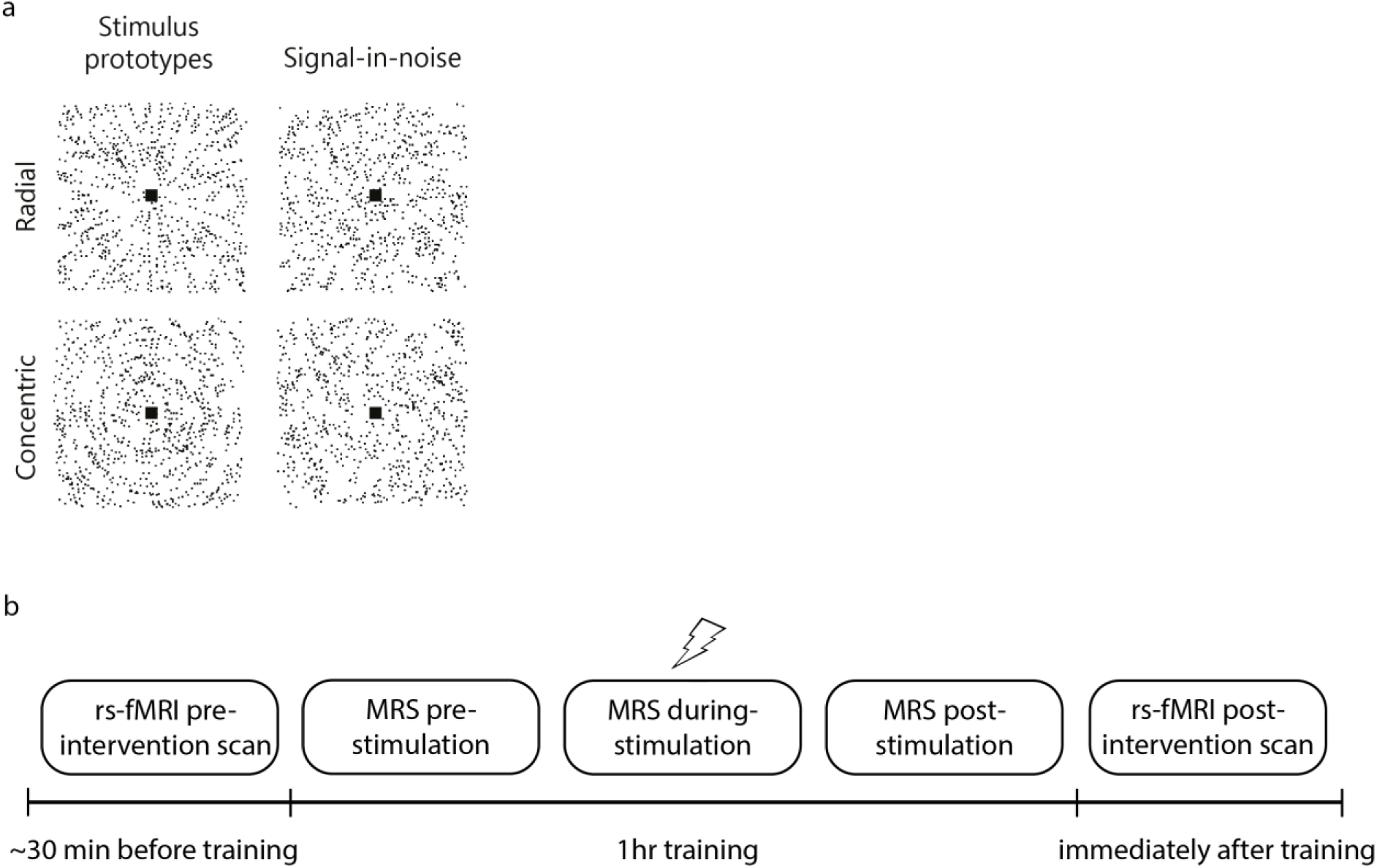
Stimuli and experiment timeline: (a) Example stimuli comprising radial and concentric Glass patterns (stimuli are presented with inverted contrast for illustration purposes). Stimuli are shown for the signal-in-noise task (25% signal, spiral angle 0° for radial and 90° for concentric). Prototype stimuli (100% signal, spiral angle 0° for radial and 90° for concentric) are shown for illustration purposes only. (b) Timeline of the experiment that comprises two rs-fMRI scans and three MRS measurements during training on the signal-in-noise task. tDCS was delivered during the second MRS acquisition for the intervention groups (Anodal, Sham).

We tested behavioural improvement in this task by comparing performance before (Pre), during (During) and after stimulation (Post) (see *Behavioural data analysis* in Methods). Our results showed that anodal OCT stimulation enhanced behavioural improvement in this task (Figure 2a), consistent with our previous work (Frangou et al., 2018). In particular, a two-way repeated measures ANOVA showed a significant Group (Anodal, Sham) x Block (Pre, During, Post) interaction (F(1.78,76.40)=3.61, p=0.037) and main effect of Block (F(1.78,76.40)=8.94, p<0.001). Performance before stimulation (i.e. Pre block) did not differ significantly between the two intervention groups (t(43)=1.02, p=0.313), suggesting that the observed differences in improvement were not due to variability in starting performance between the intervention groups. Further, comparing learning rate across training between the two groups (two-sample t-test) showed that participants in the Anodal group learned faster than participants in the Sham group (t(43)=2.31, p=0.026; Figure 2b).

**Figure 2.**
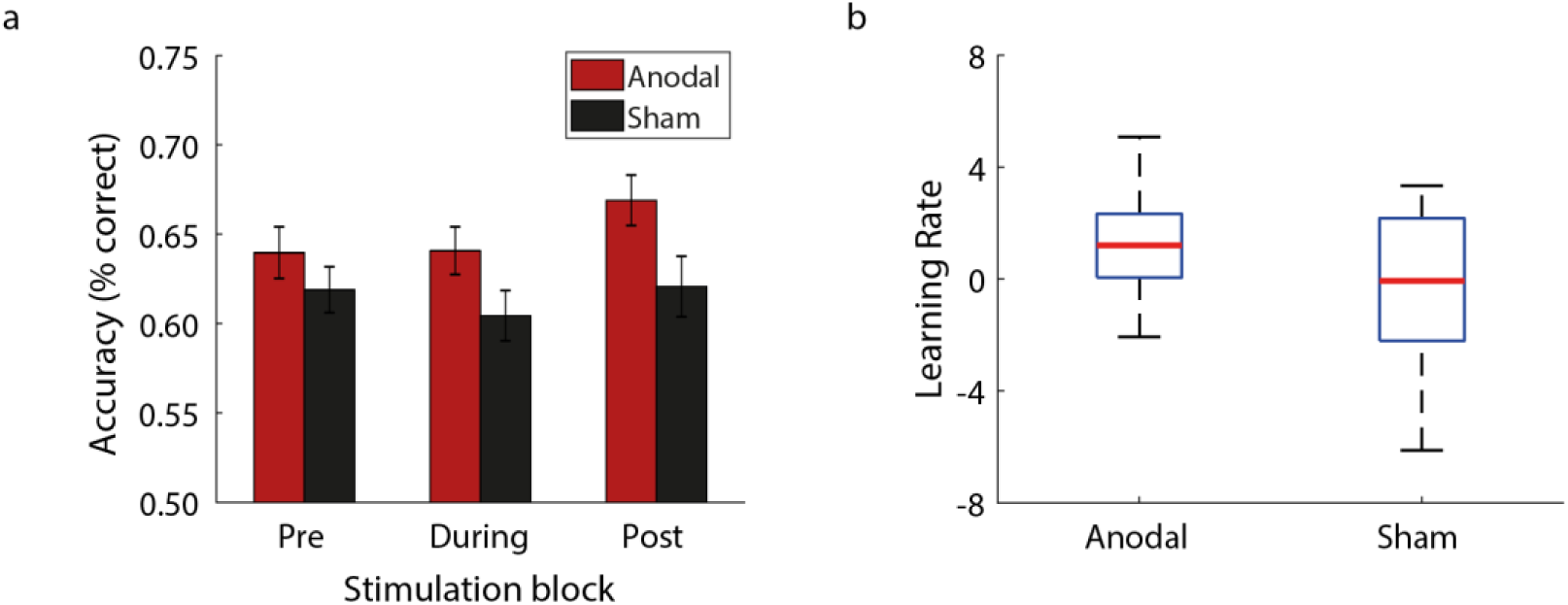
Behavioural performance: (a) Mean behavioural performance across participants per group (Anodal, Sham) and block (Pre, During, Post). Error bars indicate standard error of the mean across participants. (b) Boxplot of learning rate across training showing faster learning for Anodal than Sham group. The upper and lower error bars display the minimum and maximum data values, and the central box represents the interquartile range (25th–75th percentiles). The red line in the centre of the box represents the median.

Participants in the no-intervention (Control) group (i.e. no-stimulation, no-training group) showed no behavioural improvement in the SN task when we tested them before and after the scan (t(20)=0.18, p=0.858) nor in the contrast-detection task during the scan (one-way repeated measures ANOVA: main effect of Block: F(2.19,46.03)=0.82, p=0.458).

### Anodal tDCS results in GABA+ decrease earlier in training

To test whether anodal tDCS alters GABAergic inhibition in OCT, we measured GABA+ within an MRS voxel centred in the OCT (Figure 3) before, during and after anodal vs. sham stimulation in the OCT, while participants trained on the SN task. We compared GABA+ in the OCT for the intervention groups (i.e. anodal and sham stimulation groups who received task training) vs. the no-intervention group (i.e. no-stimulation, no-training). Comparing GABA+ change between groups, a two-way repeated measures ANOVA showed a significant Group (Anodal, Sham, Control) x Block (Pre, During, Post) interaction (F(4,120)=3.90, p=0.005; Figure 4a) and main effect of Group (F(2,60)=5.25, p=0.008). Post-hoc comparisons across blocks showed significantly decreased GABA+ for the Anodal compared to the Control group (t=-3.21, p=0.006, Bonferroni corrected), but no significant difference between Sham and Control (t=-1.93, p=0.174, Bonferroni corrected) or Anodal and Sham (t=-1.22, p=0.678, Bonferroni corrected). Further, comparing the Anodal to the Control group showed significantly decreased GABA+ for both During (t(41)=-2.23, p=0.031) and Post blocks (t(41)=-3.77, p=0.001). In contrast, comparing the Sham to the Control group showed significantly decreased GABA+ for the Post (t(40)=-2.66, p=0.011) but not the During block (t(40)=-0.88, p=0.387). These results remained significant when we tested GABA+ referenced to N acetylaspartate (NAA) rather than water (Group x Block: F(4,120)=4.06, p=0.004; main effect of Group: F(2,60)=6.35, p=0.003; Anodal vs. Control: t=-3.53, p=0.002, Bonferroni corrected; Anodal vs. Control at During block: t(41)=-2.74, p=0.009, Anodal vs. Control at Post block: t(41)=-4.46, p<0.001; Sham vs. Control at Post block: t(40)=-2.78, p=0.008). Thus, our results demonstrate that training with (anodal) or without stimulation (sham) results in decreased GABA+ in visual cortex compared to a no intervention (i.e. no training nor stimulation) control. Interestingly, training with anodal stimulation decreases GABA+ in the OCT during and after stimulation, compared to training without stimulation (i.e. sham) that shows decreases in GABA+ only after stimulation. These results suggest that anodal stimulation induces neurochemical changes earlier in the training, consistent with our behavioural results showing faster learning for anodal stimulation.

**Figure 3.**
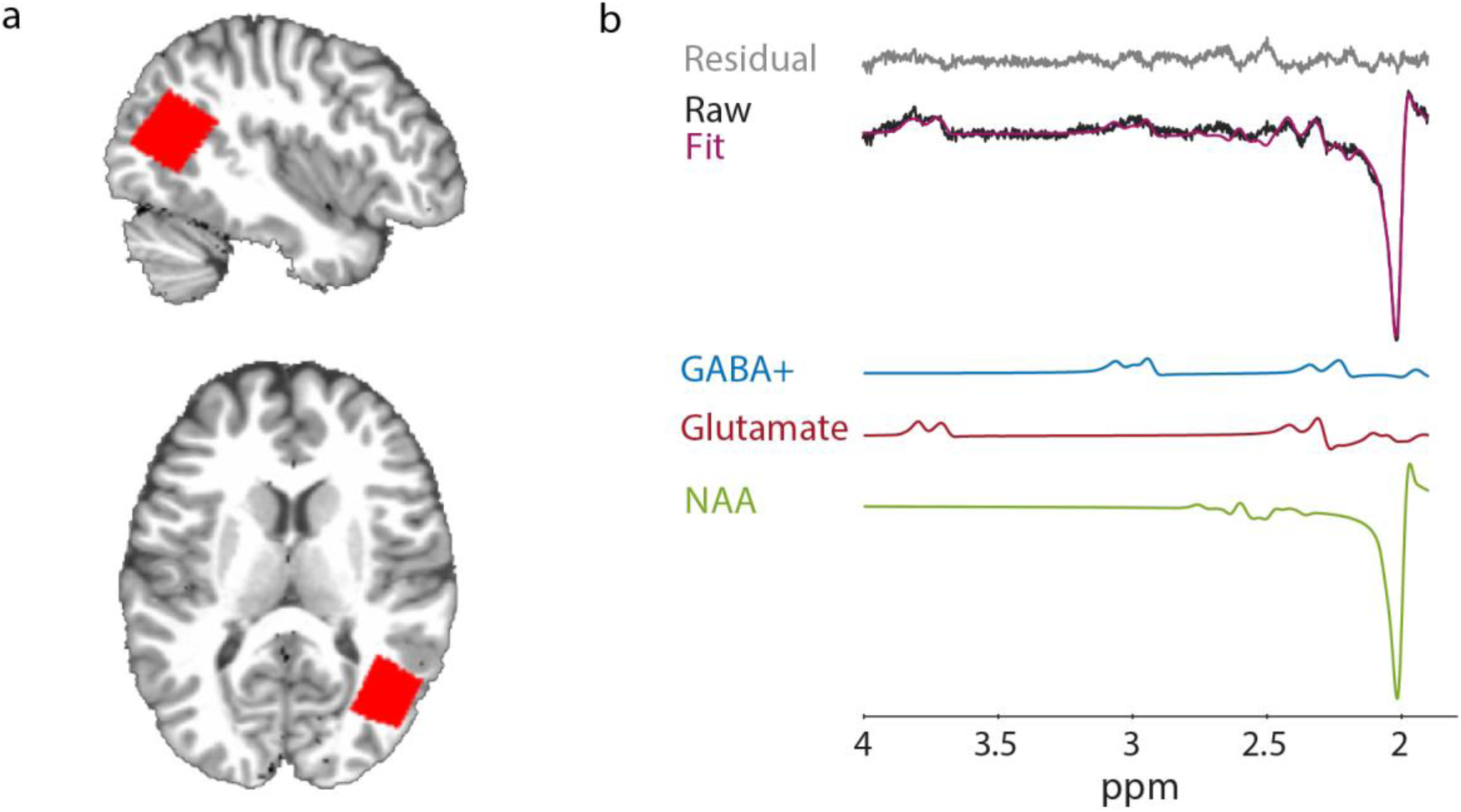
MRS voxel placement and spectra: (a) For each participant, we positioned the OCT MRS voxel using anatomical landmarks (superior temporal gyrus and middle occipital gyrus) on the acquired T1 scan to ensure that voxel placement was consistent across participants. Placement of the MRS voxel is shown for a representative participant (sagittal, axial view: native space). (b) Sample spectra from the MRS voxel of a representative participant. We show the LC model fit, the residual and the respective fits for GABA+, Glutamate and NAA.

**Figure 4.**
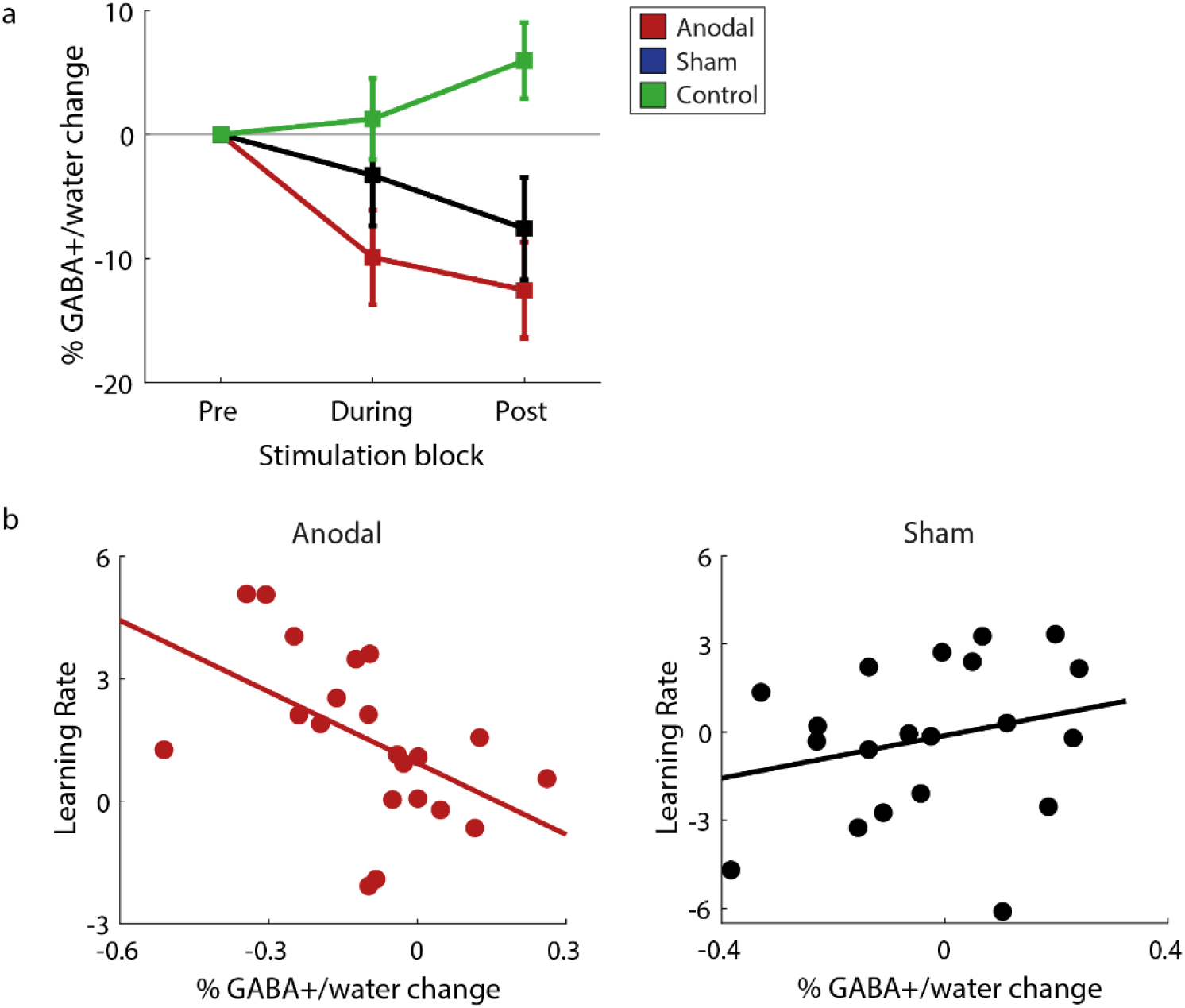
GABA+ change during training and correlations with behaviour: (a) OCT MRS-measured GABA+ over time is shown per group (Anodal, Sham, Control). We calculated % GABA+/water change by subtracting GABA+/water measurements in each of the three blocks from the pre-stimulation block and then divided by GABA+/water in the pre-stimulation block. (b) Skipped Pearson correlations showing a significant negative correlation of OCT GABA+ change (i.e. during-minus pre-stimulation block, divided by pre-stimulation block) with learning rate for the Anodal, but not the Sham group. These correlations were significantly different between groups.

It is unlikely that these changes in GABA+ for the intervention groups were due to differences in MRS data quality (i.e. linewidth, Signal-to-noise ratio: SNR) between groups (Table S1). In particular, a two-way repeated measures ANOVA showed no significant Group x Block interaction (linewidth: F(4,120)=0.32, p=0.864; SNR: F(4,120)=0.72, p=0.581) nor main effect of Group (linewidth: F(2,60)=1.44, p=0.246; SNR: F(2,60)=0.85, p=0.432). Further, GABA+ measured before stimulation did not significantly differ between groups (main effect of Group: F(2,60)=2.20, p=0.120), suggesting that our results could not be simply due to variability in GABA+ before stimulation across groups. Finally, comparing glutamate measures between groups showed no significant Group x Block interaction (F(4,120)=0.80, p=0.528) nor main effect of Group (F(2,60)=0.37, p=0.692) or Block (F(2,120)=1.33, p=0.269), suggesting that our results are specific to GABA+.

Next, we tested whether changes in OCT GABA+ relate to behavioural performance. We computed percent GABA+ change during tDCS (During) compared to GABA+ before stimulation (Pre) to control for variability in baseline GABA+ measures (i.e. Pre). We measured GABA+ change during stimulation as our previous analysis showed GABA+ changes during stimulation for the Anodal rather than the Sham group (Figure 4a). Correlating OCT GABA+ change with learning rate showed a significant negative correlation for the Anodal group (r(19)=-0.51, p=0.019; Figure 4b), but not for the Sham group (r(18)=0.25, p=0.289; Figure 4b). These correlations were significantly different between groups (Fisher’s z test: z=-2.41, p=0.016). Further, this relationship remained significant when controlling for tissue composition within the MRS voxel, controlling for MRS data quality (i.e. linewidth, SNR), and using GABA+ referenced to NAA rather than water (Table S2). There was no significant correlation for OCT Glutamate (Glu) change and learning rate, suggesting that this result is specific to GABA (Table S2). We found no significant correlation between learning rate on the contrast-detection task and change in GABA+ for the Control group (r(20)=0.06, p=0.790). Finally, there was no significant correlation between learning rate and OCT GABA+ change for the post-compared to the pre-stimulation block for any group (Anodal: r(18)=-0.19, p=0.417; Sham: r(18)=0.17, p=0.473; Control: r(20)=0.02, p=0.945), suggesting that our results are specific to the GABA+ change during stimulation. These results demonstrate that learning-dependent changes in GABA+ during training with anodal stimulation rather than training without stimulation (i.e. sham) relate to learning rate, suggesting that enhanced GABAergic plasticity due to tDCS in the OCT may facilitate faster learning in detecting targets in clutter.

### Anodal tDCS alters functional connectivity

We next tested whether anodal tDCS during training on the SN task alters extrinsic (i.e. between OCT and intra-parietal sulcus [IPS]) or intrinsic (i.e. within OCT) connectivity as measured by rs-fMRI. First, we tested for changes in extrinsic OCT-IPS connectivity after vs. before intervention. A two-way repeated measures ANOVA showed a significant Group (Anodal, Sham) x Block (Pre, Post) interaction (F(1,35)=7.96, p=0.008; Figure 5a), but no significant main effect of Group (F(1,35)=2.44, p=0.127) or Block (F(1,35)=0.01, p=0.924). Post-hoc comparisons showed a significant decrease in OCT-IPS connectivity after training for the Anodal group (t(21)=-2.16, p=0.042), but no significant change for the Sham group (t(14)=1.96, p=0.071). Next, we asked whether changes in extrinsic connectivity relate to behaviour (i.e. learning rate) and OCT GABA+ change during stimulation, as our analysis showed GABA+ changes during stimulation that relate to behaviour for the Anodal rather than the Sham group. We found a significant positive correlation of OCT-IPS connectivity change with learning rate for the Sham (r(11)=0.74, p=0.003; Figure 5b), but not for the Anodal group (r(20)=-0.24, p=0.286; Figure 5b). Comparing these correlations showed a significant difference between groups (Fisher’s z test: z=-3.06, p=0.002). Further, we found a significant positive correlation of change in OCT-IPS connectivity with change in OCT GABA+ for the Anodal (r(17)=0.48, p=0.036; Figure 5c), but not for the Sham group (r(13)=0.13, p=0.643; Figure 5c). This relationship remained significant when controlling for tissue composition within the MRS voxel, controlling for MRS data quality, and using GABA+ referenced to NAA rather than water (Table S3). There was no significant correlation for OCT Glu change and learning rate, suggesting that this result is specific to GABA (Table S3).

**Figure 5.**
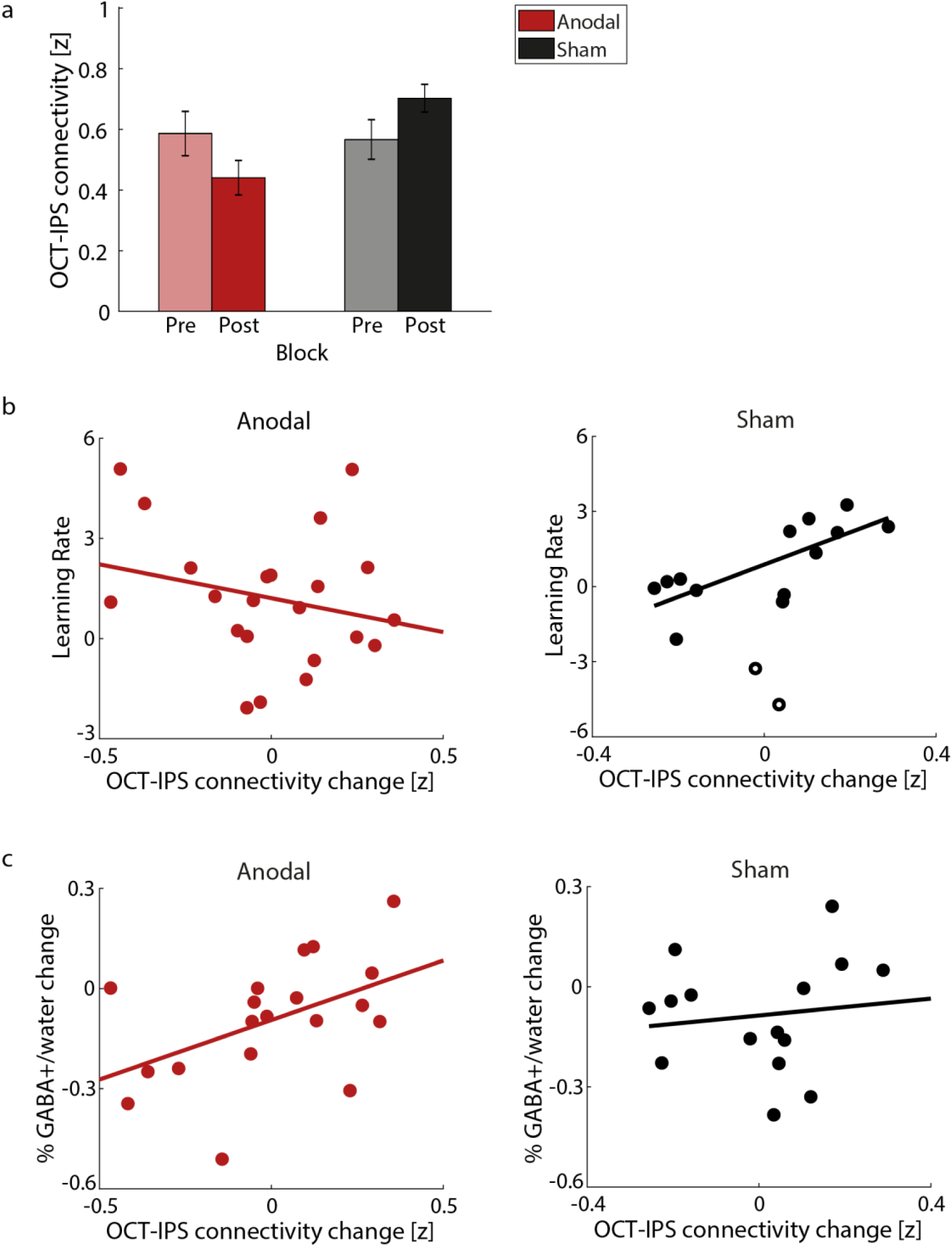
Extrinsic connectivity and correlations with behaviour and GABA+: (a) Mean OCT-IPS connectivity (Fisher z) per group (Anodal, Sham) and block (pre-, post-intervention). (b) Skipped Pearson correlations showing no significant correlation of OCT-IPS connectivity change with learning rate for the Anodal group, but a significant positive correlation for the Sham group. These correlations were significantly different between groups. (c) Skipped Pearson correlations showing a significant positive correlation of OCT-IPS connectivity change with OCT GABA+ change for the Anodal group, but not the Sham. Open symbols denote outliers.

These results demonstrate that anodal OCT stimulation results in decreased occipito-parietal connectivity after training that relates to decreased OCT GABA+ during stimulation, suggesting that enhanced GABAergic plasticity due to tDCS in the OCT may relate to local visual processing rather than occipito-parietal interactions. In contrast, for task training without stimulation (i.e. sham stimulation), learning-dependent changes in occipito-parietal connectivity relate to faster learning but not changes in GABA+.

Second, we tested for changes in intrinsic OCT connectivity after vs. before intervention. A two-way repeated measures ANOVA showed a significant main effect of Block (Pre, Post) (F(1,35)=4.66, p=0.038; Figure 6a), but no significant Group (Anodal, Sham) x Block (Pre, Post) interaction (F(1,35)=0.55, p=0.463), nor main effect of Group (F(1,35)=3.13, p=0.086). Next, we asked whether changes in intrinsic connectivity relates to learning rate and OCT GABA+ change (during-vs. pre-stimulation). There were no significant correlations of change in intrinsic OCT connectivity with learning rate (Anodal: r(20)=0.26, p=0.249; Sham: r(12)=0.52, p=0.056; Figure 6b). However, we observed a significant negative correlation for change in intrinsic OCT connectivity with change in OCT GABA+ for the Anodal group (r(14)=-0.52, p=0.039; Figure 6c), but not for the Sham group (r(13)=0.21, p=0.453; Figure 6c). This relationship remained significant when controlling for tissue composition within the MRS voxel, controlling for MRS data quality, and using GABA+ referenced to NAA rather than water (Table S3). There was no significant correlation for OCT Glu change and learning rate, suggesting that this result is specific to GABA (Table S3). Taken together, our results show that increased local OCT connectivity relates to decreases in OCT GABA+ during anodal but not sham stimulation, providing converging evidence that enhanced GABAergic plasticity due to anodal tDCS in the OCT relates to local visual processing.

**Figure 6:**
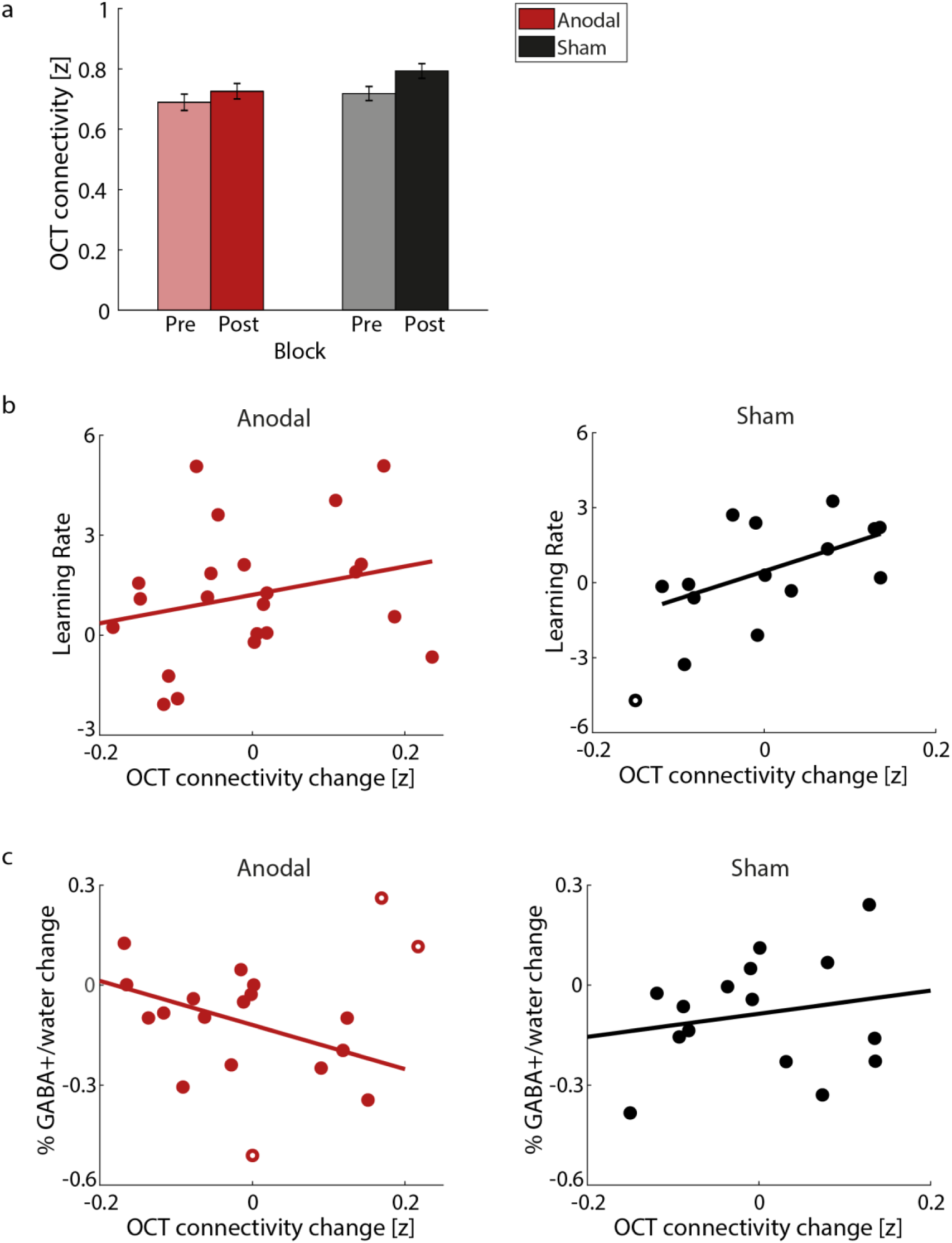
Intrinsic connectivity and correlations with behaviour and GABA+: (a) Mean intrinsic connectivity in OCT (Fisher z) per group (Anodal, Sham) and block (pre-, post-intervention). (b) Skipped Pearson correlations showing no significant correlation of OCT connectivity change with learning rate for the Anodal or the Sham group. (c) Skipped Pearson correlations showing a significant negative correlation of OCT connectivity change with OCT GABA+ change for the Anodal group, but not the Sham. Open symbols denote outliers.

### Anodal tDCS alters time-varying functional connectivity during training

Our functional connectivity analysis shows that our intervention (anodal tDCS during task training) alters occipito-parietal interactions that relate to GABAergic plasticity. However, static connectivity offers a summary measure of the synchrony between two brain regions across long timescales (i.e. 8mins for our rs-fMRI scans) that does not capture short-lived changes in inter-regional synchrony and how they propagate across different brain regions. Recent studies have proposed time-varying connectivity approaches for tracking changes in functional connectivity at finer timescales (Cohen, 2017; Hutchison et al., 2013; Preti et al., 2017). These methods have been shown to capture task and behavioural variability beyond static connectivity accounts (Calhoun et al., 2014; Eichenbaum et al., 2021). Here, we employ a time-varying connectivity analysis (i.e. HMM) to detect brain states that capture recurring patterns of activity and connectivity over time and test whether our intervention alters these brain states.

We conducted this analysis using time courses from early and higher visual areas and posterior parietal cortex as defined by a topographic atlas ((Wang et al., 2015); Table S4). We set the number of states to 5 and decomposed the input time courses to 13 Principal Component Analysis (PCA) components (corresponding to 80% variance explained) across groups and blocks. Following previous work (Karapanagiotidis et al., 2020; Vidaurre et al., 2017), we then tested the robustness of the results for a range of these parameters (states: from 4 to 7, PCA: from 70% to 100%; Table S5). The 5 estimated states capture reoccurring temporal patterns across participants and are described by a mean activation map (Figure 7a) and a functional connectivity matrix (Figure S1): State 1 captures concurrent deactivation across all regions; State 2 captures time periods when OCT (i.e. LO1/2), IPS (i.e. IPS0 and IPS1/2) and V3b are co-active; State 3 captures time periods when V1 (dorsal and ventral) V2 (dorsal) are co-active; State 4 captures concurrent activation across all regions; State 5 captures time periods when IPS (i.e. IPS0), PHC, V3a and VO are co-active. Figure 7b illustrates the transition probabilities between these states averaged across participants, where higher (lower) values represent more (less) likely transitions from one state to another.

**Figure 7:**
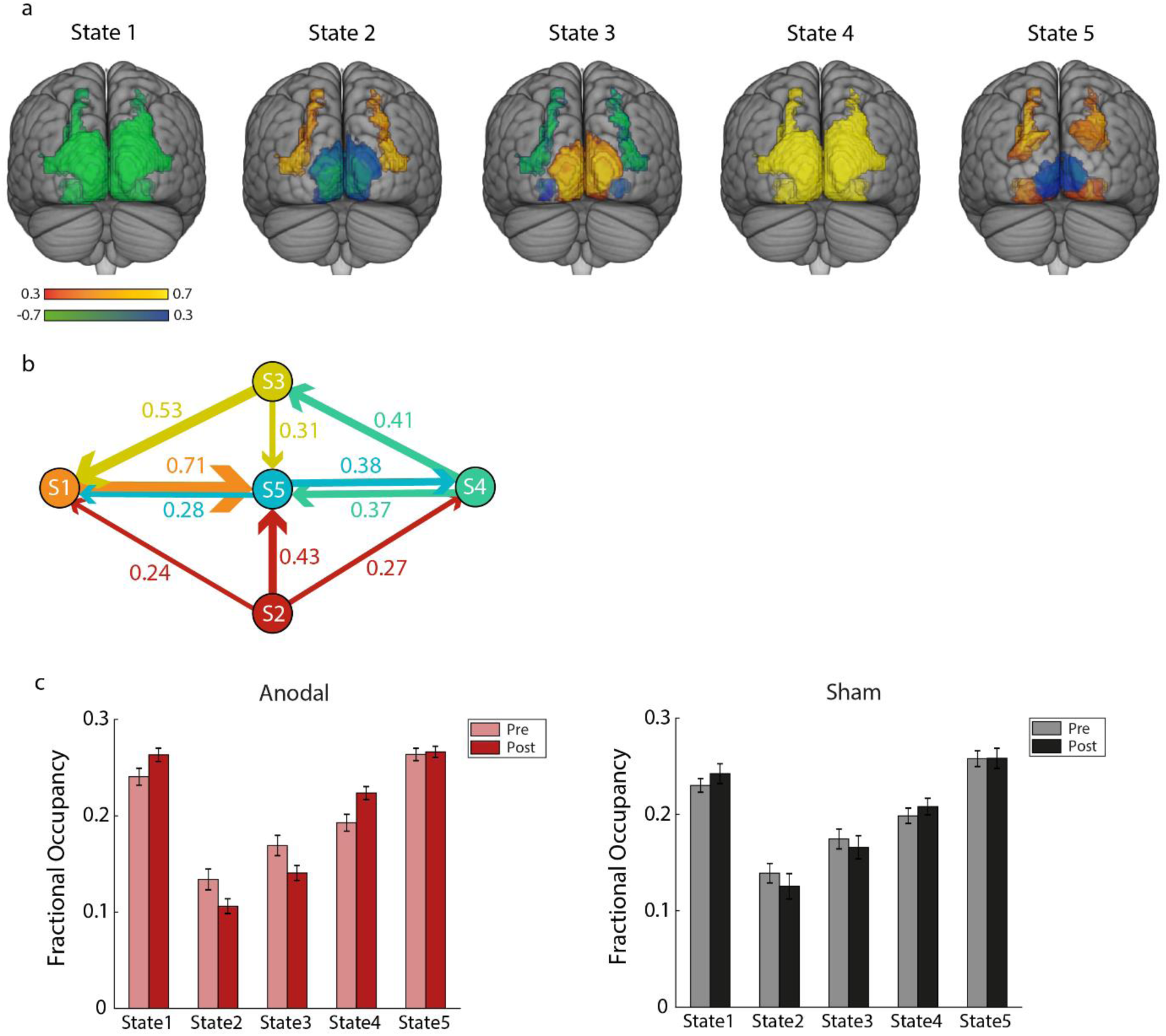
HMM states and Fractional Occupancy change: (a) Normalised mean activation maps per state are shown overlaid on the MNI brain. Warm colours (red-yellow) denote positive values, cool colours (blue-green) denote negative values for the respective region. (b) Transition probabilities between these states. The five states are displayed as nodes and the arrows denote the direction of the transition from one state to another. The thickness of the arrows is proportional to the transition probability between the corresponding states. Transition probabilities lower than 20% were removed for visualisation purposes. (c) Mean Fractional Occupancy per state and block (pre-, post-intervention) are shown for each group (Anodal: left, Sham: right). Lighter bars correspond to pre-training measures, darker bars to post-training measures.

To test for temporal differences between anodal and sham OCT stimulation, we compared the time spent in each state (i.e. Fractional Occupancy: FO) before vs. after training per group. A two-way repeated measures ANOVA showed a significant State x Block interaction for the Anodal group (F(1.45,27.93)=6.22, p=0.010; Figure 7c), but no significant effects for the Sham group (F(1.35,18.82)=1.14, p=0.319; Figure 7c). Post-hoc comparisons showed that participants in the Anodal group spent more time after training in States 1 and 4 (State 1: t(21)=2.21, p=0.038; State 4: t(21)=3.46, p=0.002); that is, they spent more time in states capturing time periods of widespread concurrent deactivation and activation across the visual cortex. Previous work has suggested that large-scale synchronised activity might denote integration of information (Varela et al., 2001) and higher sensitivity in error detection (Breakspear et al., 2003). In contrast, participants in the Anodal group spent less time in States 2 and 3 (State 2: t(21)=-2.44, p=0.023; State 3: t(21)=-2.63, p=0.015) that correspond to the OCT-IPS and the early visual (V1, V2) states, respectively. Comparing the switching rate (SR) between states before vs. after training showed that participants in the Anodal group switched more frequently between states after training (t(21)=2.30, p=0.032), while there was no significant change for the Sham group (t(14)=1.19, p=0.254).

These results complement our static connectivity results showing that anodal OCT stimulation during training in the SN task results in more widespread visual activity, and less localised activity in occipito-parietal and early visual regions. Further, anodal OCT stimulation may facilitate faster processing within brain states, as indicated by faster switching rate between states after training in the SN task.

## Discussion

Previous work has shown that training results in changes in GABAergic inhibition that relate to improved performance in visual (Frangou et al., 2019, 2018; Shibata et al., 2017) and motor tasks (Kolasinski et al., 2019; Sampaio-Baptista et al., 2015). Here, we employ tDCS to modulate cortical excitability (Antal et al., 2004a; Nitsche and Paulus, 2000) and test the role of GABAergic inhibition in perceptual learning. tDCS has been shown to alter performance in visual tasks (Antal et al., 2004b; Battaglini et al., 2017; Spiegel et al., 2013; Zito et al., 2015) and facilitate learning in motor (O’Shea et al., 2017) and visual memory tasks (Barron et al., 2016) by reducing GABA. Here, we demonstrate that modulating GABAergic inhibition with tDCS during training boosts performance in perceptual judgements by altering local processing in visual cortex and functional connectivity between visual and posterior parietal areas that are involved in perceptual decision making. Our findings advance our understanding of GABAergic plasticity mechanisms for perceptual learning in the following respects.

First, we have previously shown that anodal tDCS during training enhances behavioural improvement on the signal-in-noise task that has been shown to relate to decreased GABAergic inhibition (Frangou et al., 2018). Here, we demonstrate that tDCS dissociates faster vs. slower learning and GABAergic plasticity. In particular, we show that training with anodal stimulation on the visual cortex results not only in behavioural improvement, but also faster learning compared to training without stimulation (i.e. sham stimulation). Further, training with anodal tDCS results in decreased OCT GABA+ during and after stimulation, in contrast to training without stimulation (i.e. sham) that shows a later decrease in OCT GABA+ (i.e. after stimulation). Next, we show that this decrease in GABA+ during anodal stimulation relates to faster learning, suggesting that anodal OCT stimulation during training accelerates perceptual learning by shifting GABAergic plasticity in the OCT earlier in the learning process.

Second, previous studies have shown that training in a range of tasks (e.g. motor or perceptual tasks) results in changes in functional connectivity (Guerra-Carrillo et al., 2014; Karlaftis et al., 2019; Kelly and Castellanos, 2014; Lewis et al., 2009; Sampaio-Baptista et al., 2015). Further, functional connectivity has been shown to relate to GABAergic inhibition (Frangou et al., 2019; Kapogiannis et al., 2013; Karlaftis et al., 2021; Mann and Paulsen, 2007; Nasrallah et al., 2017; Sampaio-Baptista et al., 2015; Shmuel and Leopold, 2008; Stagg et al., 2014) and can be altered by tDCS during training in a range of tasks: spatial navigation (Krishnamurthy et al., 2015), associative learning (Krause et al., 2017), language processing (Cao and Liu, 2018; Meinzer et al., 2012), visual selective attention (McDermott et al., 2019) and visual search (Callan et al., 2016). In our previous work, we showed that perceptual learning in the signal-in-noise task relates to functional connectivity within visual cortex and between visual and posterior parietal regions measured at rest before training (Frangou et al., 2019). Here, we test whether combining brain stimulation with training results in changes in functional connectivity. Our results demonstrate that anodal —rather than sham— OCT stimulation during training, decreases occipito-parietal connectivity. This is consistent with previous work showing that IPS is involved in identifying salient and task-relevant features early in training, while OCT is involved in tuning task-relevant feature representation after training (Chang et al., 2014; Frangou et al., 2019; Mayhew et al., 2012). In particular, our results show that increased occipito-parietal connectivity relates to faster learning for participants in the Sham group, who show slower improvement and are therefore engaged in earlier stages of learning. In contrast, for the Anodal group, occipito-parietal connectivity shows a significant decrease that correlates with OCT GABA+ decrease during our intervention (stimulation and training). The relationship between tDCS-induced changes in GABAergic inhibition and functional connectivity remains debated, with some studies showing a significant relationship (Antonenko et al., 2017), but some others not (e.g. (Bachtiar et al., 2015)). Here, we show that anodal tDCS during training to detect targets in clutter results in accelerated GABA decrease in visual cortex that relates to reduced occipito-parietal connectivity, suggesting that anodal tDCS alters functional connectivity between sensory and decision-related areas.

Further, we show a significant negative correlation between intrinsic connectivity change and OCT GABA+ change during anodal but not sham stimulation (potentially due to the delayed GABA decrease in the Sham group). This relationship between changes in OCT GABA+ and local temporal coherence suggests that decreased GABAergic inhibition within visual cortex may facilitate signal detection by enhancing local processing. This result is consistent with our previous work showing higher intrinsic connectivity before training for greater GABA decrease during training (Frangou et al., 2019).

Previous work has shown that time-varying connectivity captures task and behavioural variability in addition to what is explained by static connectivity (Calhoun et al., 2014; Liégeois et al., 2019; Vidaurre et al., 2021). Here, we employ HMM to detect brain states of recurrent activity and connectivity patterns that have been linked to cognition (Karapanagiotidis et al., 2020; Vidaurre et al., 2017) and investigate learning-dependent plasticity at a finer timescale. We show that anodal stimulation alters inter-regional synchrony at both coarse (i.e. static functional connectivity over longer time periods, in the range of minutes) and finer timescales (i.e. functional changes within shorter time windows, in the range of seconds). In particular, we find decreased localised activity (occipito-parietal, early visual) after training with anodal stimulation, consistent with the decreased static occipito-parietal connectivity. In contrast, we find increased widespread synchronised activation across the whole visual cortex after training with anodal stimulation. Widespread synchronised activity has been linked to integration of information (Varela et al., 2001) and higher sensitivity in error detection (Breakspear et al., 2003). These processes are key for our signal-in-noise task that involves integrating information across space, detecting the relevant features (i.e. signal) and suppressing irrelevant information (i.e. noise).

Finally, despite the wide interest that tDCS has attracted in cognitive and clinical neuroscience, its validity remains debated and our understanding of the tDCS mechanisms of action remains limited (Fertonani and Miniussi, 2017). Here we address this challenge by combining tDCS with brain imaging to interrogate the brain mechanisms that underlie the facilitatory effect of tDCS on learning and brain plasticity. Our findings dissociate faster vs. slower learning mechanisms and provide evidence for GABAergic plasticity mechanisms across stages of learning. In particular, we demonstrate that tDCS results in faster learning to detect targets in clutter by accelerating GABAergic plasticity (i.e. reducing GABAergic inhibition) and decreasing occipito-parietal connectivity. Our findings propose that brain stimulation during training optimises sensory processing through local gain control mechanisms (i.e. reduction of GABAergic inhibition) (Katzner et al., 2011) to support improved perceptual decisions (i.e. detecting targets in cluttered scenes).

## Materials and Methods

### Participants

We tested forty-five healthy volunteers (27 female; mean age 22.9 ± 3.3 years) in two intervention groups, twenty-four in the stimulation group (Anodal) and twenty-one in the no-stimulation group (Sham). We tested an additional no-intervention group of twenty-two healthy volunteers who did not receive training nor stimulation (Control: 17 female; mean age 25.8 ± 4.2 years). All participants were right-handed, had normal or corrected-to-normal vision, did not receive any prescription medication, were naïve to the aim of the study, gave written informed consent and received payment for their participation. The study was approved by the University of Cambridge Ethics Committee [PRE.2017.057].

### Stimuli

We presented participants with Glass patterns (Glass, 1969) generated using previously described methods ((Zhang et al., 2010); Figure 1a). In particular, stimuli were defined by white dot pairs (dipoles) displayed within a square aperture on a black background. Stimuli (size=7.9° x 7.9°), were presented in the left hemifield (11.6 arc min from fixation) contralateral to the stimulation site to ensure maximal effect of stimulation on stimulus processing. The dot density was 3%, and the Glass shift (i.e., the distance between two dots in a dipole) was 16.2 arc min. The size of each dot was 2.3 x 2.3 arc min^2^. For each dot dipole, the spiral angle was defined as the angle between the dot dipole orientation and the radius from the centre of the dipole to the centre of the stimulus aperture. Each stimulus comprised dot dipoles that were aligned according to the specified spiral angle (signal dipoles) for a given stimulus and noise dipoles for which the spiral angle was randomly selected. The proportion of signal dipoles defined the stimulus signal level.

We generated radial (0° spiral angle) and concentric (90° spiral angle) Glass patterns by placing dipoles orthogonally (radial stimuli) or tangentially (concentric stimuli) to the circumference of a circle centred on the fixation dot. A new pattern was generated for each stimulus presented in a trial, resulting in stimuli that were locally jittered in their position. Radial (spiral angle: 0°) and concentric stimuli (spiral angle: ± 90°) were presented at 23% or 25% signal level counterbalanced across trials; noise dipoles were presented at random position and orientation. To control for potential local adaptation due to stimulus repetition and ensure that learning related to global shape rather than local stimulus features, we jittered (± 1-3°) the spiral angle across stimuli.

### Experimental Design

All participants in the intervention groups took part in a single brain imaging session during which they were randomly assigned to the Anodal or Sham group. Participants in the Anodal group received anodal tDCS on the right OCT, whereas participants in the Sham group did not receive stimulation. We recorded three MRS measurements from the right OCT during training: before, during and after stimulation. In addition, we recorded whole-brain rs-fMRI data before and after training while participants fixated on a cross at the centre of the screen (Figure 1b). Participants in the no-intervention Control group took part in a single brain imaging session without stimulation or training; we recorded three MRS measurements from right OCT at the same timings of the MRS measurements as for the intervention groups. We did not record rs-fMRI data for this group due to time constraints.

During training, participants in the intervention groups were presented with Glass patterns and were asked to judge and indicate by button press whether the presented stimulus in each trial was radial or concentric. Two stimulus conditions (radial vs. concentric Glass patterns; 100 trials per condition), were presented for each training block. For each trial, a stimulus was presented for 300ms and was followed by fixation (i.e., blank screen with a central fixation dot) while waiting for the participant’s response (self-paced training paradigm). Trial-by-trial feedback was provided by means of a visual cue (green tick for correct, red ‘x’ for incorrect) followed by a fixation dot for 500ms before the onset of the next trial.

In the no-intervention control group, participants were tested in a contrast change detection task. In particular, participants were presented with Glass patterns where 100% of the dipoles were randomly oriented (0% signal patterns). In each trial, participants were asked to choose whether the top or bottom half of the pattern underwent a contrast change. Task difficulty was controlled by a two-up-one-down staircase to ensure participants were not trained at the task and response accuracy was held at 75%.

### MRI data acquisition

We collected MRI data on a 3T Siemens PRISMA scanner (Cognition and Brain Sciences Unit, Cambridge) using a 64-channel head coil. T1-weighted structural data (TR = 19.17s; TE = 2.31ms; number of slices = 176; voxel size = 1mm isotropic) and echo-planar imaging (EPI) data (gradient echo-pulse sequences) were acquired during rest (TR = 0.727s; TE = 34.6ms; number of slices = 72; voxel size = 2mm isotropic; Multi-band factor = 8; flip angle = 51°; number of volumes = 660; duration = 8m09s; whole brain coverage). During EPI data acquisition, we recorded cardiac pulsation (using a pulse oximeter) and respiration (using a respiratory belt) to model these physiological data for denoising.

### MRS data acquisition

We collected MRS data with a MEGA-PRESS sequence (Mescher et al., 1998): echo time = 68ms, repetition time = 3000ms; 256 transients of 2048 data points were acquired in 13min experiment time; a 14.28ms Gaussian editing pulse was applied at 1.9 (ON) and 7.5 (OFF) ppm; water unsuppressed 16 transients (Table S6, following guidelines by (Lin et al., 2021)). Measurements with this sequence at 3T have been previously shown to produce reliable and reproducible estimates of GABA+ (Puts and Edden, 2012). Water suppression was achieved using variable power with optimized relaxation delays and outer volume suppression. Automated shimming was conducted to achieve water linewidth below 10Hz. We acquired spectra from an MRS voxel (20 x 20 x 25 mm^3^) in the right OCT (Figure 3). We manually positioned the MRS voxel using anatomical landmarks (superior temporal gyrus, middle occipital gyrus) on each participant’s structural scan, ensuring that voxel placement was consistent across participants. The centre of gravity for the MRS voxel was: x=40.8±3.2mm, y=-61.7±5.2mm, z=10.6±3.6mm in Montreal Neurological Institute (MNI) space. During the MRS acquisitions, participants in the intervention groups performed the SN task, while participants in the no-intervention control group performed the contrast change detection task.

### tDCS data acquisition

We used a multi-channel transcranial electrical stimulator (neuroConn DC-STIMULATOR MC, Ilmenau, Germany) to deliver anodal or sham stimulation. We used a pair of MR-compatible rubber electrodes (3x3 cm^2^ stimulating electrode, 5x5 cm^2^ reference electrode), which were secured on the head with the help of rubber bands. Ten-20 paste was used as a conductive medium between the rubber electrodes and the scalp. For the Anodal group, 1mA current was ramped up over 10s, was held at 1mA for 20min and was subsequently ramped down over 10s. For the Sham group, the current ramped up (10s) and down (10s) in the beginning of the session. We used online stimulation (i.e. stimulation during training), as this protocol has been previously shown to enhance the lasting effect of training (O’Shea et al., 2017). It has been shown that this facilitatory effect is not present or polarity-specific when stimulation precedes training, with anodal stimulation impeding learning (Stagg et al., 2011). To achieve consistent electrode placement across participants when targeting the right posterior OCT (consistent with the MRS acquisition in the right OCT), we placed the bottom right corner of the square stimulating electrode on T6, using a 10-20 system EEG cap, maintaining the same orientation across participants, parallel to the line connecting T6 and O2. The reference electrode was placed on Cz. We have previously used the same electrode montage (Frangou et al, 2018), following electrical field density simulations showing that this montage results in unilaterally localised current density, the peak of the electric field density being under the anode electrode around the posterior OCT and the stimulation reaching the region where the MRS voxel was placed.

### Behavioural data analysis

We measured behavioural performance per training block as the mean accuracy per 200 trials. To quantify learning-dependent changes in behaviour, we computed the behavioural performance before, during and after stimulation as the average performance of blocks 1-2 (Pre), 3-5 (During) and 6-9 (Post), respectively. Further, we quantified learning rate by fitting individual participant training data with a logarithmic function: y = *k* ∗ ln *x* + c, where *x* is the training run separated into 100 trial bins, *y* is the run accuracy, *c* is the starting performance and *k* corresponds to the learning rate. Positive learning rate indicates that performance improved with training, whereas negative or close to zero learning rate indicates no behavioural improvement.

### MRS data analysis

We pre-processed the MRS data using MRspa v1.5c (www.cmrr.umn.edu/downloads/mrspa/). We applied Eddy current, frequency and phase correction before subtracting the average ON and OFF spectra, resulting in edited spectra. We used LC-Model (Provencher, 2001) to quantify metabolite concentrations by fitting simulated model spectra of γ-amino-butyric acid (GABA), Glu, Glutamine and NAA to the edited spectra (Figure 3b), setting the sptype parameter to mega-press-2. We refer to GABA concentration as GABA+, as MRS measurements of GABA with MEGA-PRESS include co-edited macromolecules (Mullins et al., 2014). We referenced metabolite concentrations to the concentration of water for our analyses and then validated our findings by referencing GABA+ to NAA to ensure our results were not driven by the chosen reference (Lunghi et al., 2015).

GABA+ measurements within 3 standard deviations from the mean across all groups and blocks (data for 1 participant of the Anodal group were excluded) and with Cramer-Rao lower bound (CRLB) values smaller than 15% (data for 2 participants of the Anodal group and 1 of the Sham group were excluded) were included in further steps of MRS related analyses. SNR was calculated as the amplitude of the NAA peak in the difference-spectrum divided by twice the root mean square of the residual signal (Provencher, 2001). We report average quality indices (CRLB, linewidth, SNR) per group and block (Table S1). To control for potential differences in data quality across participants and blocks, we performed control analyses that accounted for changes in linewidth and SNR (Table S2, Table S3). We did not include control analyses for changes in CRLB, as reductions in GABA concentration have been shown to be inherently linked to increases in CRLB (Emir et al., 2012; Kreis, 2016; Lunghi et al., 2015).

Further, we conducted whole brain tissue-type segmentation of the T1-weighted structural scan and calculated percentage of grey matter, white matter and cerebrospinal fluid (CSF) in the MRS voxel. To ensure that correlations with GABA+ were not driven by variability in tissue composition within the MRS voxel across participants, we conducted two control analyses (Table S2, Table S3): (a) regressed out the CSF percentage from the GABA+ concentrations, (b) applied α-correction on the GABA+ values to account for the difference in GABA+ between grey and white matter (Porges et al., 2017).

### rs-fMRI data analysis

We pre-processed the structural and the rs-fMRI data in SPM12.4 (v7219; www.fil.ion.ucl.ac.uk/spm/software/spm12/) following the Human Connectome Project pipeline for multi-band data (Smith et al., 2013). In particular, we first coregistered (non-linearly) the T1w structural images (after brain extraction) to MNI space to ensure that all participant data were in the same stereotactic space for statistical analysis. We then (a) corrected the EPI data for susceptibility distortions (fieldmap correction) and any spatial misalignments between EPI volumes due to head movement (i.e. aligned each run to its single band reference image), (b) coregistered the second EPI run to the first (rigid body) to correct any spatial misalignments between runs, (c) coregistered the first EPI run to the structural image (rigid body) and (d) normalised them to MNI space for subsequent statistical analyses (applying the deformation field of the structural images). Data were only resliced after MNI normalisation to minimise the number of interpolation steps. Following MNI normalisation, (e) data were skull-stripped, (f) spatially smoothed with a 4mm Gaussian kernel to improve the signal-to-noise ratio and the alignment between participant data (two times the voxel size; (Chen and Calhoun, 2018)), (g) wavelet despiked to remove any secondary motion artifacts (Patel et al., 2014) and (h) had linear drifts removed (linear detrending due to scanner noise). Slice-timing correction was not applied, following previous work on fast TR (sub-second) acquisition protocols (Smith et al., 2013). Data from 8 participants (2 anodal, 6 sham) were excluded from further analysis due to missing the second rs-fMRI run.

Next, we applied spatial group Independent Component Analysis (ICA) using the Group ICA fMRI Toolbox (GIFT v3.0b) (http://mialab.mrn.org/software/gift/) to identify and remove components of noise. PCA was applied for dimensionality reduction, first at the subject level, then at the group level. The Minimum Description Length criteria (Rissanen, 1978) were used to estimate the dimensionality and determine the number of independent components. The ICA estimation (Infomax) was run 20 times and the component stability was estimated using ICASSO (Himberg et al., 2004). Following recent work on back-reconstruction methods for ICA denoising at the group level (Du et al., 2016), we used Group Information Guided ICA (GIG-ICA) back-reconstruction to reconstruct subject-specific components from the group components. We visually inspected the results and identified noise components according to published procedures (Griffanti et al., 2017). Using consensus voting among 3 experts, we labelled 8 of the 31 components as noise that captured signal from veins, arteries, CSF pulsation, susceptibility and multi-band artefacts.

To clean the fMRI signals from signals related to motion and the noise components, we followed a soft clean-up ICA denoise approach (Griffanti et al., 2014). That is, we first regressed out the motion parameters (translation, rotation and their squares and derivatives; (Friston et al., 1996)) from each voxel and ICA component time course. Second, we estimated the contribution of every ICA component to each voxel’s time course (multiple regression). Finally, we subtracted the unique contribution of the noise components from each voxel’s time course to avoid removing any shared signal between neuronal and noise components.

Following ICA denoise, we performed a first-level analysis modelling the physiological data as nuisance variables. We used the TAPAS toolbox (Kasper et al., 2017) to create physiological covariates that model terms for RETROICOR (Glover et al., 2000), heart rate variability (Chang et al., 2009) and respiratory volume per time (Birn et al., 2008). Following previous work (Caballero-Gaudes and Reynolds, 2017), we selected a second-order model for both the cardiac and the respiratory signal (no interaction term) and zero delay for the heart rate variability and respiratory volume per time terms. Within the GLM, the data were high-pass filtered at 0.01Hz and treated for serial correlations using the FAST autoregressive model, as it has been shown to perform more accurate autocorrelation modelling for fast TR acquisitions (Corbin et al., 2018; Olszowy et al., 2019). The residual time course from the last step was used for all subsequent analyses.

### Static connectivity analysis

We calculated extrinsic functional connectivity between OCT and IPS and intrinsic connectivity within OCT. First, we created masks for these two regions of interest (ROI). For OCT, we computed the overlap across participant MRS voxels and created a group MRS mask that included voxels present in at least 50% of the participants’ MRS voxels. For IPS, we created an equally sized cubic mask centred on the intraparietal cortex (centre at 34, -50, 42 in MNI space (Frangou et al., 2019), edge length = 20mm).

Then, for each participant and ROI, we computed the first eigenvariate across all grey matter voxels within the region to derive a single representative time course per ROI. We applied a 5th order Butterworth band-pass filter between 0.01 and 0.08 Hz on the eigenvariate time course, similar to previous studies (Cordes et al., 2001; Frangou et al., 2019; Murphy et al., 2013). Extrinsic functional connectivity was computed as the Pearson correlation of the OCT-IPS time courses. Similarly, intrinsic connectivity was computed as the Pearson correlation of each OCT voxel’s time course to the eigenvariate time course and then averaged across voxels (Bachtiar et al., 2015; Frangou et al., 2019; Stagg et al., 2014; Van Dijk et al., 2010). We computed the change in rs-fMRI connectivity as the difference of the pre-from the post-intervention run (after Fisher z-transform) and tested for: (a) changes in extrinsic and intrinsic connectivity, (b) correlations of connectivity change with OCT GABA+ change, and (c) correlations of connectivity change with behaviour. For correlations with GABA+ and behaviour, we regressed out the pre-intervention connectivity from the difference to control for baseline differences across participants.

### Time-varying connectivity analysis

We estimated time-varying functional connectivity using the HMM-MAR toolbox (Vidaurre et al., 2018, 2017). In particular, we estimated a HMM on the visual cortex to detect brain states representing recurrent patterns of activity and connectivity over time. Using a Bayesian approach, the model learns a set of parameters for each state and the probability of their activation at each time point given the recorded data. Specifically, given an active state *Z*_*t*_ at time *t*, the recorded data sample *X*_*t*_ is described by a multivariate Gaussian distribution: *P*(*X*_*t*_|*Z*_*t*_ = *k*) ∼ *N*(*μ*_*k*_, Σ_*k*_). Each state has distinct mean and covariance parameters that capture each state’s mean activation and functional connectivity.

To investigate the dynamics of the occipito-parietal (OCT-IPS) interactions with the rest of the visual cortex, we defined fourteen bilateral regions from the Probabilistic map of Visual Topography (Wang et al., 2015) (Table S4). We then computed the first eigenvariate across all voxels within each region to derive a single representative time course per ROI. We concatenated the time courses of all ROIs across participants and runs to estimate state distributions (i.e. the spatial parameters of the model) at the group level, whereas the probability of a state activation is still defined uniquely for each timepoint at the participant level (i.e. the temporal parameters of the model; (Vidaurre et al., 2016)).

Latent variable models (such as the HMM) can be sensitive to local minima or poor initialisation (Vidaurre et al., 2019). To ensure stability on the estimation of the HMM states, we ran the algorithm 10 times with 10 random initialisations for each iteration and selected the iteration with the lowest free energy for simplicity. Further, we tested whether the results were robust to variations of key hyperparameters: the number of states ranging from 4 to 10, and the input data dimensionality by varying the number of retained PCA dimensions to capture between 70% and 100% of the variance (in increments of 10%).

To describe the state dynamics, we computed two summary measures: FO per state, as the proportion of time spent in that state, and SR across states, as the frequency of switching between states. That is, a state with increased (decreased) FO after training indicates that regions within that state are more (less) involved in the processing of the task, suggesting a higher (lower) engagement of that state after training. Similarly, increased SR after training indicates faster switching from one state to another over time, suggesting shorter processing times within a state after training. Finally, we computed change in FO and SR as the difference of the pre-from the post-intervention rs-fMRI run and tested for within-group changes.

### Statistical analysis

For ANOVAs, we tested for sphericity and used Greenhouse-Geisser (for epsilon less than 0.75) or Huynh-Feldt (for epsilon greater than 0.75) correction, if sphericity was violated. For correlational analyses, we used skipped Pearson correlation of the Robust Correlation Toolbox to account for bivariate outliers and adjusted the degrees of freedom when outliers were detected (Pernet et al., 2013).

### Funding

This work was supported by grants to ZK from the Biotechnology and Biological Sciences Research Council (H012508, BB/P021255/1), the Wellcome Trust (205067/Z/16/Z) and the European Community’s Seventh Framework Programme (FP7/2007-2013) under agreement PITN-GA-2011-290011. For the purpose of open access, the author has applied for a CC BY public copyright licence to any Author Accepted Manuscript version arising from this submission.

## Acknowledgements

We would like to thank the MR physics and radiographer teams at the Cognition and Brain Sciences Unit for their support with data collection, and Matthew Davis and Benedikt Zoefel for their guidance on setting up tDCS in the scanner. We would like to thank Vicki Hodgson for help with data collection and Joseph Giorgio for his help with reviewing the ICA components.

## Abbreviations

CRLB: Cramer-Rao lower bound
CSF: cerebrospinal fluid
EPI: echo-planar imaging
FO: Fractional Occupancy
Glu: Glutamate
HMM: Hidden Markov Models
ICA: Independent Component Analysis
IPS: intra-parietal sulcus
MNI: Montreal Neurological Institute
MRS: Magnetic Resonance Spectroscopy
NAA: N acetylaspartate
OCT: occipito-temporal cortex
PCA: Principal Component Analysis
ROI: region of interest
rs-fMRI: resting-state functional Magnetic Resonance Imaging
SN: signal-in-noise
SNR: Signal-to-noise ratio
SR: Switching Rate
tDCS: transcranial direct current stimulation

---

**Figure S1:**
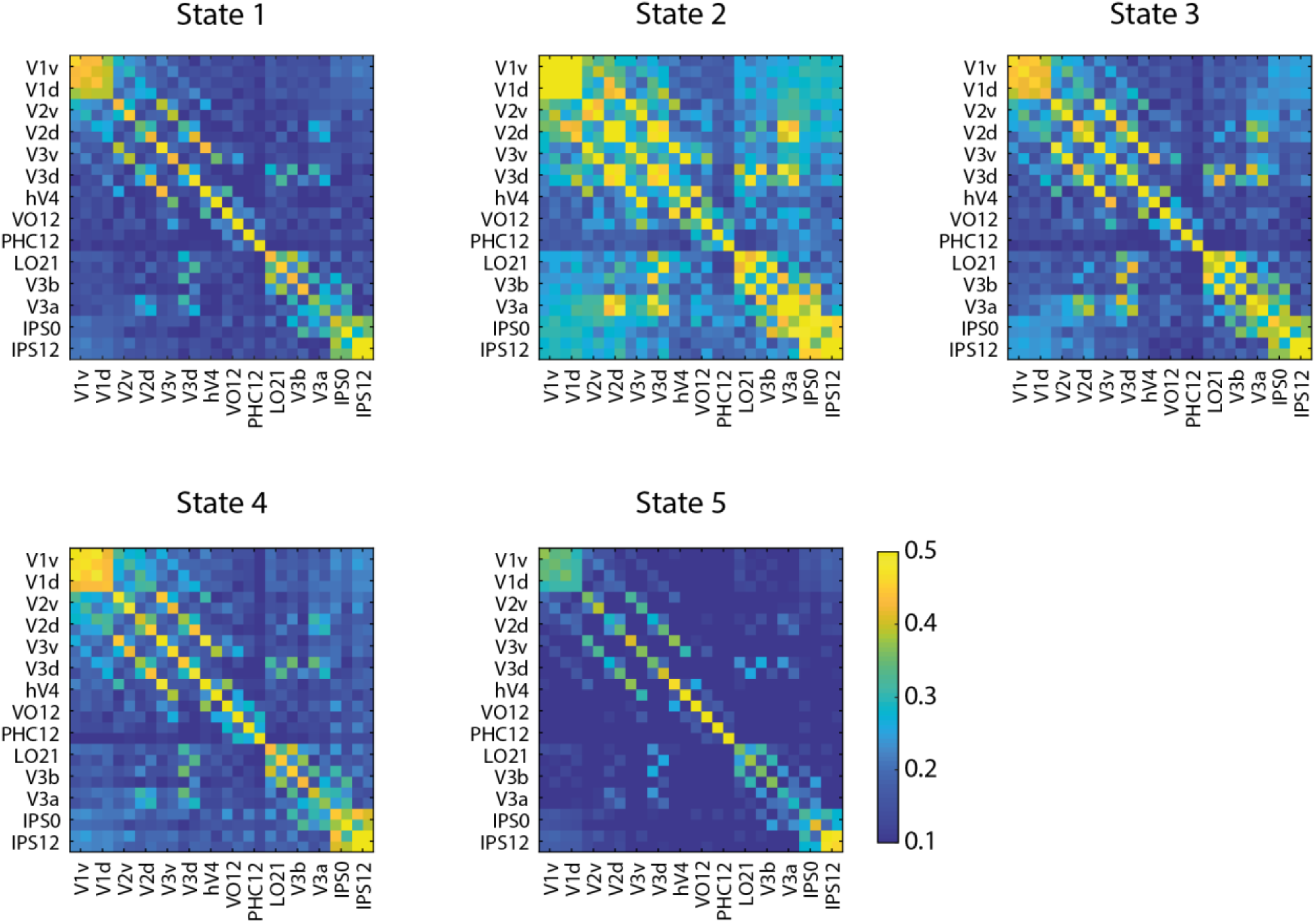
Functional Connectivity matrices of HMM states: Functional Connectivity matrices are shown as 28x28 matrices per state. For each ROI, data are included for each hemisphere (left, right). Warm colours (yellow) denote higher connectivity values, cool colours (blue) denote connectivity values close to zero.

## Supplementary Tables

**Table S1.** MRS quality measures: Cramer-Rao lower bound (CRLB), linewidth and signal-to-noise ratio (SNR) are shown for the OCT MRS voxel per group and block.

| MRS quality measure | Block | Group | Mean | Std |
| --- | --- | --- | --- | --- |
| **CRLB** | Pre | Anodal | 5.67 | 1.20 |
|  |  | Sham | 5.65 | 1.14 |
|  |  | Control | 5.91 | 1.31 |
|  | During | Anodal | 6.76 | 2.00 |
|  |  | Sham | 6.35 | 2.13 |
|  |  | Control | 6.23 | 1.27 |
|  | Post | Anodal | 6.81 | 1.89 |
|  |  | Sham | 6.50 | 1.73 |
|  |  | Control | 5.82 | 1.05 |
| **Linewidth** | Pre | Anodal | 7.83 | 0.67 |
|  |  | Sham | 7.96 | 0.45 |
|  |  | Control | 8.14 | 0.48 |
|  | During | Anodal | 7.94 | 0.70 |
|  |  | Sham | 8.07 | 0.50 |
|  |  | Control | 8.20 | 0.53 |
|  | Post | Anodal | 7.95 | 0.68 |
|  |  | Sham | 8.15 | 0.49 |
|  |  | Control | 8.22 | 0.56 |
| **SNR** | Pre | Anodal | 19.95 | 3.40 |
|  |  | Sham | 20.50 | 2.98 |
|  |  | Control | 19.27 | 3.58 |
|  | During | Anodal | 19.33 | 3.50 |
|  |  | Sham | 20.00 | 3.34 |
|  |  | Control | 18.68 | 3.63 |
|  | Post | Anodal | 20.29 | 3.77 |
|  |  | Sham | 20.15 | 3.01 |
|  |  | Control | 18.91 | 3.44 |

**Table S2:** Control analyses for GABA+ correlation with behaviour: Pearson correlations (r, p) for GABA+ change and learning rate when: regressing out the CSF percentage or using α-correction to control for tissue composition within the MRS mask, using GABA+ referenced to NAA (rather than water), regressing out changes in MRS data quality (linewidth, SNR), and testing for neurotransmitter specificity (Glu change).

| Group | Control | r | p |
| --- | --- | --- | --- |
| **Anodal** | %CSF | -0.52 | 0.017 |
| | α-correction | -0.51 | 0.018 |
|  | GABA+/NAA | -0.47 | 0.031 |
|  | Linewidth | -0.49 | 0.024 |
|  | SNR | -0.50 | 0.022 |
|  | Glu | -0.28 | 0.225 |
| **Sham** | %CSF | 0.25 | 0.291 |
| | α-correction | 0.26 | 0.262 |
|  | GABA+/NAA | 0.27 | 0.248 |
|  | Linewidth | 0.25 | 0.291 |
|  | SNR | 0.27 | 0.251 |
|  | Glu | -0.07 | 0.782 |

**Table S3:** Control analyses for GABA+ correlation with resting-state connectivity: Pearson correlations (r, p) for GABA+ change and resting-state connectivity when: regressing out the CSF percentage or using α-correction to control for tissue composition within the MRS mask, using GABA+ referenced to NAA (rather than water), regressing out changes in MRS data quality (linewidth, SNR), and testing for neurotransmitter specificity (Glu change).

| Connectivity | Group | Control | r | p |
| --- | --- | --- | --- | --- |
| **OCT-IPS** | **Anodal** | %CSF | 0.48 | 0.039 |
| | | α-correction | 0.50 | 0.030 |
|  |  | GABA+/NAA | 0.51 | 0.025 |
|  |  | Linewidth | 0.49 | 0.032 |
|  |  | SNR | 0.48 | 0.037 |
|  |  | Glu | 0.03 | 0.909 |
|  | **Sham** | %CSF | 0.24 | 0.396 |
| | | α-correction | 0.12 | 0.667 |
|  |  | GABA+/NAA | 0.19 | 0.495 |
|  |  | Linewidth | 0.14 | 0.620 |
|  |  | SNR | 0.13 | 0.656 |
|  |  | Glu | -0.22 | 0.426 |
| **OCT** | **Anodal** | %CSF | -0.50 | 0.048 |
| | | α-correction | -0.53 | 0.035 |
|  |  | GABA+/NAA | -0.54 | 0.033 |
|  |  | Linewidth | -0.52 | 0.037 |
|  |  | SNR | -0.51 | 0.045 |
|  |  | Glu | 0.08 | 0.773 |
|  | **Sham** | %CSF | 0.29 | 0.299 |
| | | α-correction | 0.20 | 0.466 |
|  |  | GABA+/NAA | 0.25 | 0.377 |
|  |  | Linewidth | 0.26 | 0.358 |
|  |  | SNR | 0.26 | 0.353 |
|  |  | Glu | -0.13 | 0.633 |

**Table S4.** Visual regions for time-varying connectivity analysis: Regions were selected from the Probabilistic map of Visual Topography (Wang et al., 2015). The size and the MNI coordinates of the centre of gravity for each region are shown. Regions between 30 and 100 voxels were grouped together with a neighbouring region that serves similar functionality and displayed a similar time course. Regions smaller than 30 voxels were excluded from the analysis as signals being unreliable.

| Region | Hem. | Size | x | y | z |
| --- | --- | --- | --- | --- | --- |
| V1v | L | 462 | -6 | -89 | -5 |
|  | R | 394 | 8 | -87 | -2 |
| V1d | L | 389 | -7 | -96 | 2 |
|  | R | 359 | 11 | -94 | 5 |
| V2v | L | 359 | -9 | -83 | -11 |
|  | R | 368 | 10 | -81 | -8 |
| V2d | L | 309 | -10 | -99 | 12 |
|  | R | 336 | 15 | -96 | 14 |
| V3v | L | 247 | -17 | -79 | -12 |
|  | R | 280 | 18 | -77 | -11 |
| V3d | L | 264 | -18 | -97 | 16 |
|  | R | 253 | 24 | -94 | 16 |
| hV4 | L | 155 | -25 | -80 | -14 |
|  | R | 173 | 26 | -79 | -12 |
| VO1, VO2 | L | 192 | -25 | -66 | -10 |
|  | R | 214 | 26 | -64 | -9 |
| PHC1, PHC2 | L | 176 | -27 | -52 | -8 |
|  | R | 164 | 28 | -49 | -9 |
| LO1, LO2 | L | 228 | -35 | -88 | 7 |
|  | R | 221 | 38 | -85 | 9 |
| V3b | L | 111 | -29 | -90 | 17 |
|  | R | 152 | 34 | -84 | 18 |
| V3a | L | 195 | -19 | -91 | 24 |
|  | R | 359 | 21 | -88 | 28 |
| IPS0 | L | 264 | -25 | -80 | 30 |
|  | R | 235 | 30 | -78 | 33 |
| IPS1, IPS2 | L | 206 | -21 | -71 | 46 |
|  | R | 155 | 25 | -69 | 47 |

**Table S5:** Control analyses for time-varying connectivity analysis: Repeated measures ANOVA results (State x Block interaction on Fractional Occupancy) are shown per group for a range of HMM parameters (states: from 4 to 7, PCA: from 70% to 100% in increments of 10%). F and p-values are reported per test and the significant results are shown in italic.

| **Model** | **Group** | **F** | **p** |
| --- | --- | --- | --- |
| states=4, PCA=70% | *Anodal* | *7.36* | *0.006* |
|  | Sham | 0.62 | 0.486 |
| states=4, PCA=80% | *Anodal* | *7.81* | *0.006* |
|  | Sham | 1.12 | 0.318 |
| states=4, PCA=90% | *Anodal* | *7.42* | *0.009* |
|  | Sham | 1.10 | 0.318 |
| states=4, PCA=100% | *Anodal* | *9.70* | *0.004* |
|  | Sham | 1.73 | 0.210 |
| states=5, PCA=70% | *Anodal* | *5.55* | *0.013* |
|  | Sham | 0.74 | 0.451 |
| states=5, PCA=80% | *Anodal* | *6.22* | *0.010* |
|  | Sham | 1.14 | 0.319 |
| states=5, PCA=90% | *Anodal* | *7.50* | *0.007* |
|  | Sham | 0.66 | 0.474 |
| states=5, PCA=100% | *Anodal* | *7.59* | *0.005* |
|  | Sham | 1.84 | 0.194 |
| states=6, PCA=70% | *Anodal* | *5.99* | *0.005* |
|  | Sham | 0.41 | 0.688 |
| states=6, PCA=80% | *Anodal* | *6.84* | *0.005* |
|  | Sham | 0.52 | 0.598 |
| states=6, PCA=90% | *Anodal* | *7.23* | *0.008* |
|  | Sham | 0.54 | 0.536 |
| states=6, PCA=100% | *Anodal* | *6.93* | *0.009* |
|  | Sham | 1.87 | 0.190 |
| states=7, PCA=70% | *Anodal* | *4.38* | *0.013* |
|  | Sham | 0.47 | 0.688 |
| states=7, PCA=80% | *Anodal* | *5.40* | *0.008* |
|  | Sham | 0.65 | 0.556 |
| states=7, PCA=90% | *Anodal* | *6.52* | *0.008* |
|  | Sham | 0.63 | 0.521 |
| states=7, PCA=100% | *Anodal* | *4.84* | *0.009* |
|  | Sham | 1.54 | 0.237 |

**Table S6:**
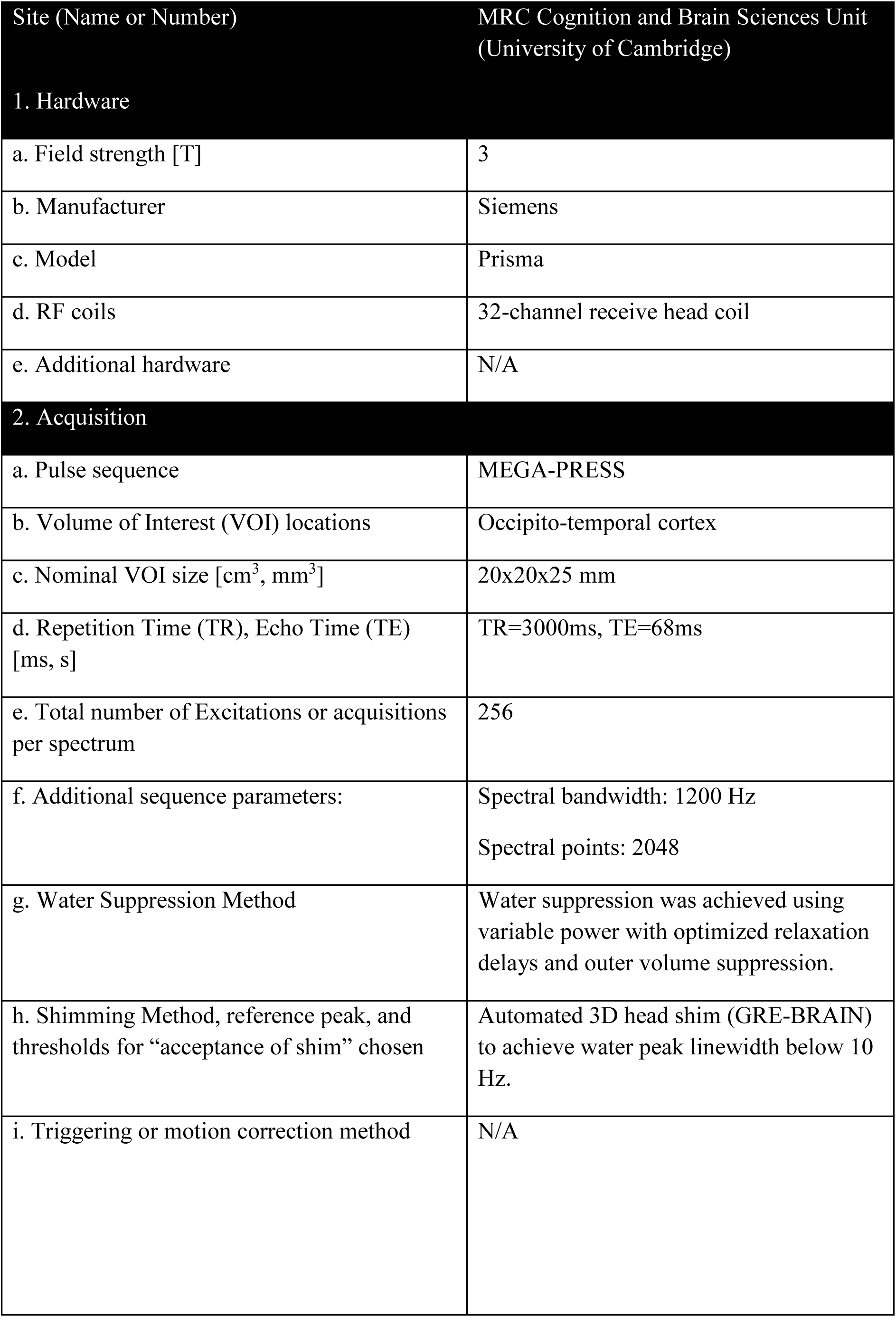
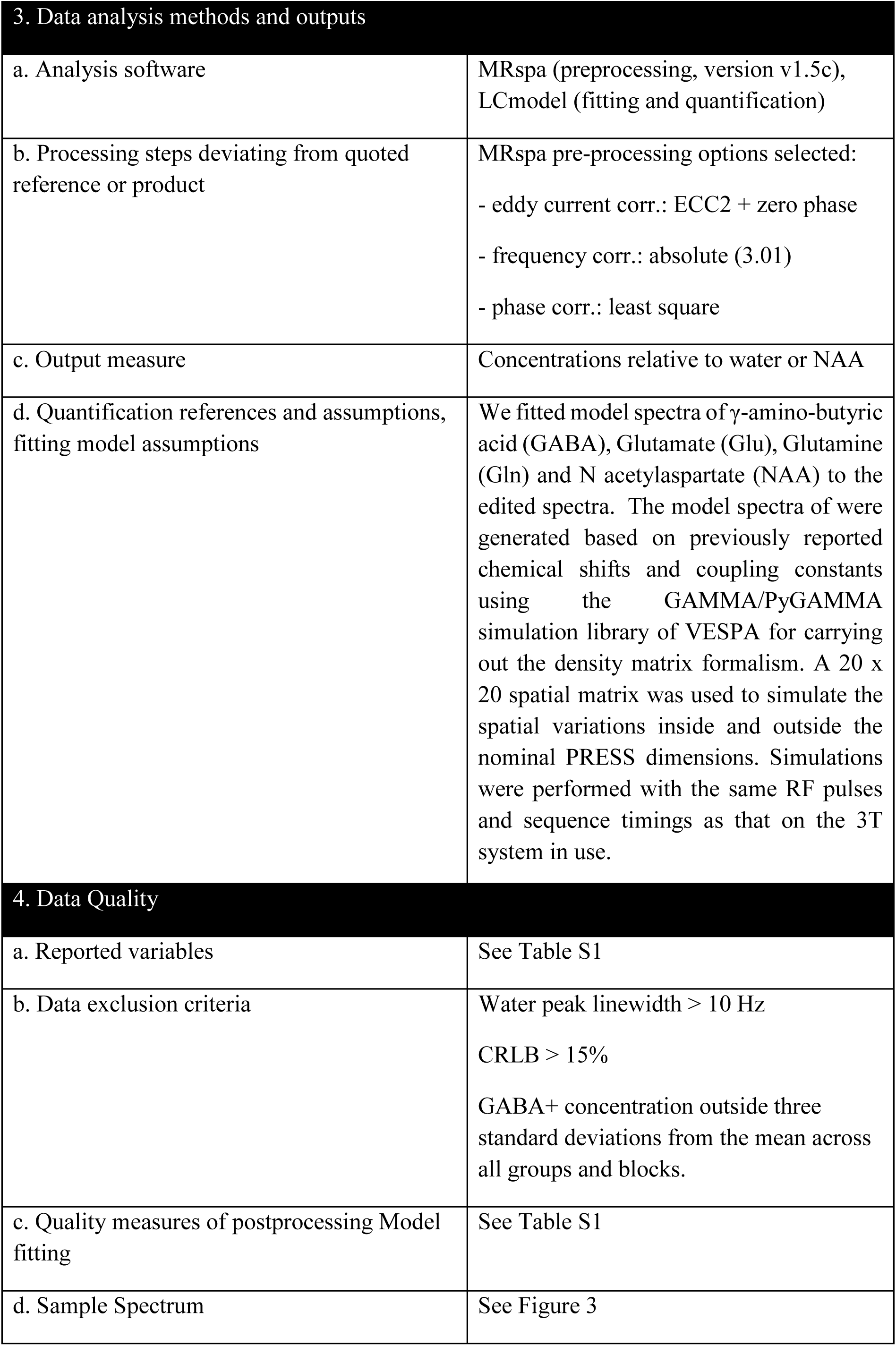
Minimum Reporting Standards in MRS checklist.

